# Evolutionary contingency facilitates scent divergence in an emergent orchid population

**DOI:** 10.64898/2026.09.26.753903

**Authors:** Jasen W. Liu, Kathy Darragh, Maria F. Guizar Amador, Dawson Mai, Cheryl A. Dean, Diego Bogarín, Santiago R. Ramírez

## Abstract

Reproductive isolation can evolve rapidly even among populations with extensive genetic similarity, but the mechanisms underlying this divergence remain an open question in evolutionary biology. Here, we investigate the genetic basis of floral scent divergence, a key trait mediating differential pollinator attraction, in a group of nascent *Gongora* orchid lineages that occur in sympatry. We combined whole-genome resequencing and floral transcriptomics to characterize the transition from the monoterpene-emitting “M chemotype” to the pollinator-isolated phenylpropanoid-emitting “A chemotype” nested within it. While overall genetic differentiation was low, a few highly differentiated regions contained candidate genes in biosynthetic pathways. Of particular interest was a putative o-methyltransferase exhibiting evidence of a recent selective sweep in the A chemotype, with much higher expression in scent-emitting floral tissues of A than in M. Transcriptomic comparison with a more distantly related monoterpene-emitting lineage (“X chemotype”) further revealed the coordinated and elevated expression of phenylpropanoid biosynthetic genes in M despite its absence of aromatic floral volatiles, including a putative key enzyme potentially involved in directing metabolic flux towards monolignol biosynthesis. Together, our results support a model in which pre-existing activity in a metabolic pathway may facilitate the evolution of a novel floral scent phenotype, potentially making transitions across the adaptive landscape of scent more accessible than others. We propose that such evolutionary contingency may help explain how divergent scent profiles, and consequently, reproductive isolation, can evolve rapidly via few genetic changes.

## Introduction

Signaling traits that contribute to reproductive isolation often evolve rapidly during speciation ^1^. In animal-pollinated flowering plants, floral scent often functions as a sexual signal by mediating pollinator attraction and successful pollen transfer among conspecific individuals ^2,3^. Therefore, divergence in floral scent between populations can result in the attraction of pollinators with distinct olfactory preferences, a process that may reduce gene flow and facilitate the formation of species boundaries ^4–6^. Floral scent is highly evolutionarily labile, with its rapid evolution attributed to the diversification of several groups exhibiting specialized pollination systems ^7,8^. Despite the importance of floral scent in shaping fundamental evolutionary processes, understanding how this complex trait diverges so rapidly has received relatively little attention.

Perfume flowers offer a unique opportunity to understand the mechanistic basis of scent divergence in the context of pollinator isolation. Floral scent emission in these species both acts as an attractant for their male euglossine bee pollinators as well as well as a reward as they are collected and incorporated into species-specific blends of perfume used during courtship ^9,10^. As such, scent determines pollinator identity, and shifts in these chemical phenotypes can result in pollinator divergence and consequently, reproductive isolation. Scent evolves rapidly in perfume flowers, with frequent shifts observed between biosynthetic pathways (phenylpropanoid and terpenoid) at both coarse and fine scales of biological organization^11^. The apparent ease of these shifts is thought to drive diversification across perfume flowers, resulting in higher speciation rates associated with this pollination mode^7,12^. Thus, determining the genetic basis of such shifts thus provides valuable insight into the mechanisms underlying the rapid radiation of a group of flowering plants.

Orchids of the genus *Gongora* offer an excellent model system to begin investigating these questions. This genus is one of the most diverse genera of perfume flowers, containing 60-70 recognized species found in lowland to mid-elevation rainforests from Mexico to Argentina^13^. Taxonomy of the group has been hindered by the occurrence of cryptic species complexes, where multiple “chemotypes” co-occur in a region that exhibit nearly identical morphological traits with radically different floral chemistries ^14–16^. Guizar Amador (2022) used an integrative approach to investigate four sympatric chemotypes in the Golfo Dulce region of Costa Rica, using a combination of population genetics and pollinator observations to characterize their divergent scent profiles and pollinator visitation patterns concomitant with subtle genetic differentiation (Fig. 1A). In particular, one of the chemotypes produces a monoterpene-dominated “piney” scent, consisting of mostly the cyclic monoterpene terpinen-4-ol (the “M” chemotype), while a lineage nested within it produces a slightly “fishy” or “eggy” scent dominated by the volatile aromatic compounds estragole and *cis*- and *trans*- isomers of methyl methoxy cinnamate (the “A” chemotype). These chemotypes both morphologically resemble *G. gibba* (G. Gerlach, personal communication) and exhibit overlapping color phenotypes. The strong differences in scent are associated with pollination by almost completely non-overlapping assemblages of male *Euglossa* bees, making the A-M comparison a compelling system to understand rapid phenotypic divergence in chemistry and pollinator isolation.

**Figure 1.**
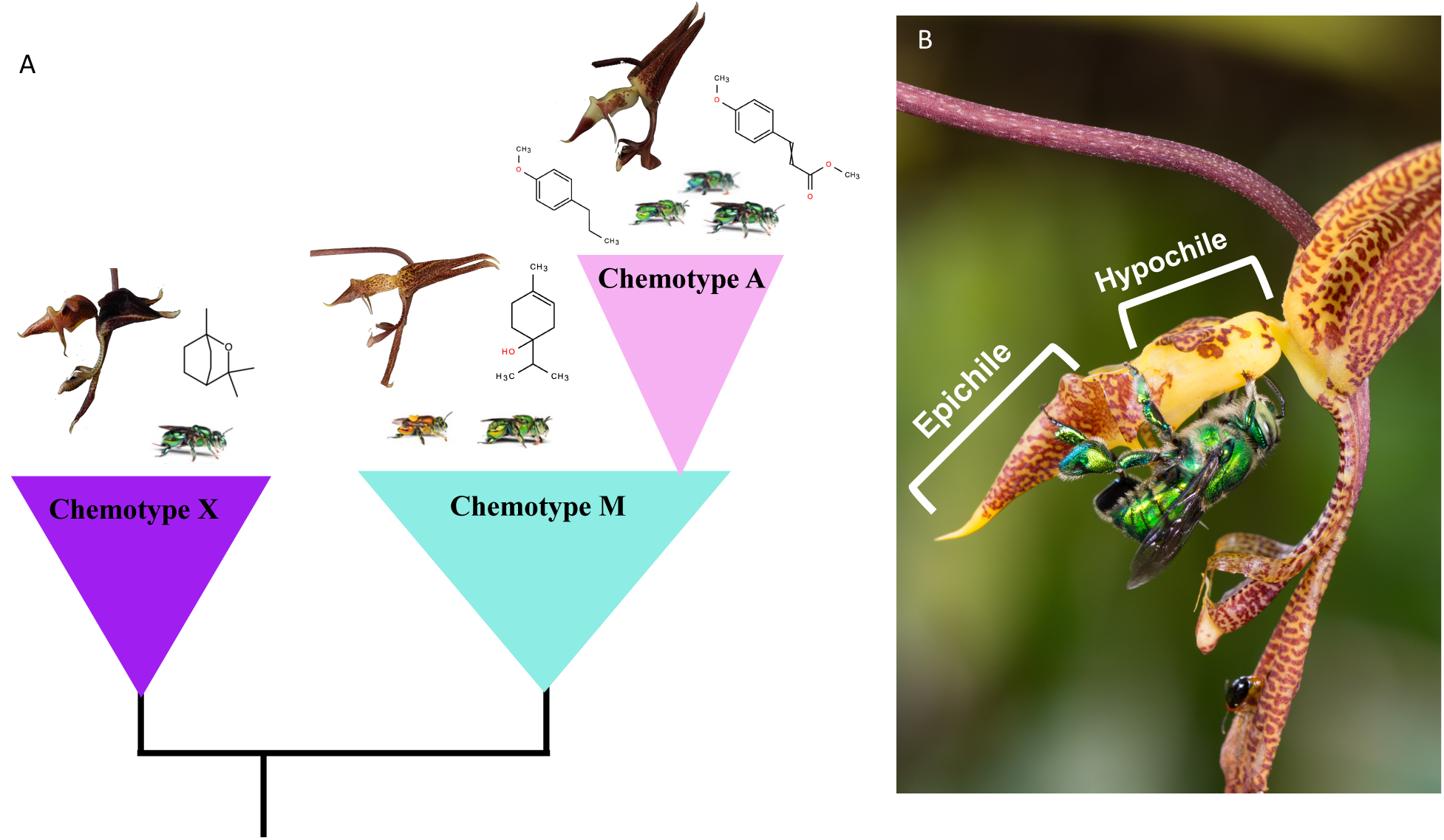
Natural history of the study system. A) Phylogenetic framework modified from Guizar Amador (2022) of three sympatric chemotypes of *Gongora* orchids growing in the Golfo Dulce Region of Costa Rica. Line drawings of chemicals represent the dominant compounds in the scent of the three chemotypes (from left to right, 1,8-cineole, terpinen-4-ol, estragole, and methyl methoxycinnamate). Images of bees represent dominant pollinators (*Euglossa imperialis* in X; *E. hansoni* and *tridentata* in M; *E. variabilis*, *E. milenae*, and *E. imperialis* in A; photographs taken by T. Eltz). B) Scent-collecting behavior on a *Gongora* flower by a *Euglossa* male, where the bee collects scent compounds emitted from the hypochile region on the labellum. Foraging in this position places him such that if he slips, he lands on the reproductive organs directly below. Photograph taken by T. Eltz.

The contrasting chemistries of these lineages are underlain by distinct biosynthetic mechanisms with an extensive history of molecular biology research in plants. Isoprenoid precursors to the monoterpenes produced by the M chemotype are synthesized via the methylerythritol phosphate (MEP) pathway and are subsequently catalyzed by terpene synthases (TPSs) into their final products ^17^. Small changes in TPS coding sequences can result in divergent products, and thus this class of enzymes is thought to be a major driver of diversity across this large chemical class ^18^. By contrast, the aromatic compounds produced by the A chemotype are phenylpropenes, volatile derivatives of phenylpropanoid biosynthesis^19,20^. Divergence between scent profiles likely results from recruitment of enzymes within these pathways between closely-related species, directing metabolic flux towards the production of different substrates. Pre-existing transcriptional activity leading to an altered metabolic background could thus provide a new source of genetic variation for selection to act upon, allowing disproportionately large phenotypic changes to arise through relatively few genetic modifications ^21,22^. Consequently, such pre-existing activity (“latent metabolic potential”) could facilitate certain transitions across the adaptive landscape of floral scent, particularly when the resulting phenotypes alter pollinator visitation.

Here, we combine whole-genome resequencing and floral transcriptomics to investigate the genetic basis of rapid scent divergence between the pollinator-isolated A and M chemotypes. We first performed genome-wide scans of F_ST_ and identified divergence concentrated in a relatively small number of differentiated peaks. We then performed an RNA-sequencing experiment, finding substantial tissue-specific differential expression of several genes found in these genetically differentiated regions that were putatively involved in scent production. Finally, we compared gene expression of the M chemotype with another more distantly related monoterpene-dominated chemotype (the “X” chemotype; Fig. 1A), and identified upregulation of many genes associated with phenylpropanoid biosynthesis, suggesting an altered metabolic background predisposing it to phenotypic transition in volatiles. Our integrative analyses provide valuable insight into understanding how floral scent, a key trait mediating pollinator isolation, evolves so rapidly.

## Results

### Scent divergence occurs despite limited genome-wide differentiation

We performed whole genome sequencing of 10 A and 10 M individuals and generated an average of 148.5 million reads per sample, with over 99% mapping rate across all samples resulting in an average of 14.5x coverage and no significant difference between chemotypes (Fig. S1A). The chemotypes clustered clearly in multivariate space, with the first principal components of genetic variation based on 290,366 unlinked sites clearly differentiating the two (Fig. 2A). However, further axes of variation showed no separation of the chemotypes (Fig. 2B). We further identified low genetic differentiation between the chemotypes, with a genome-wide F_ST_ weighted average across 7,124,345 SNPs of 0.035.

**Figure 2.**
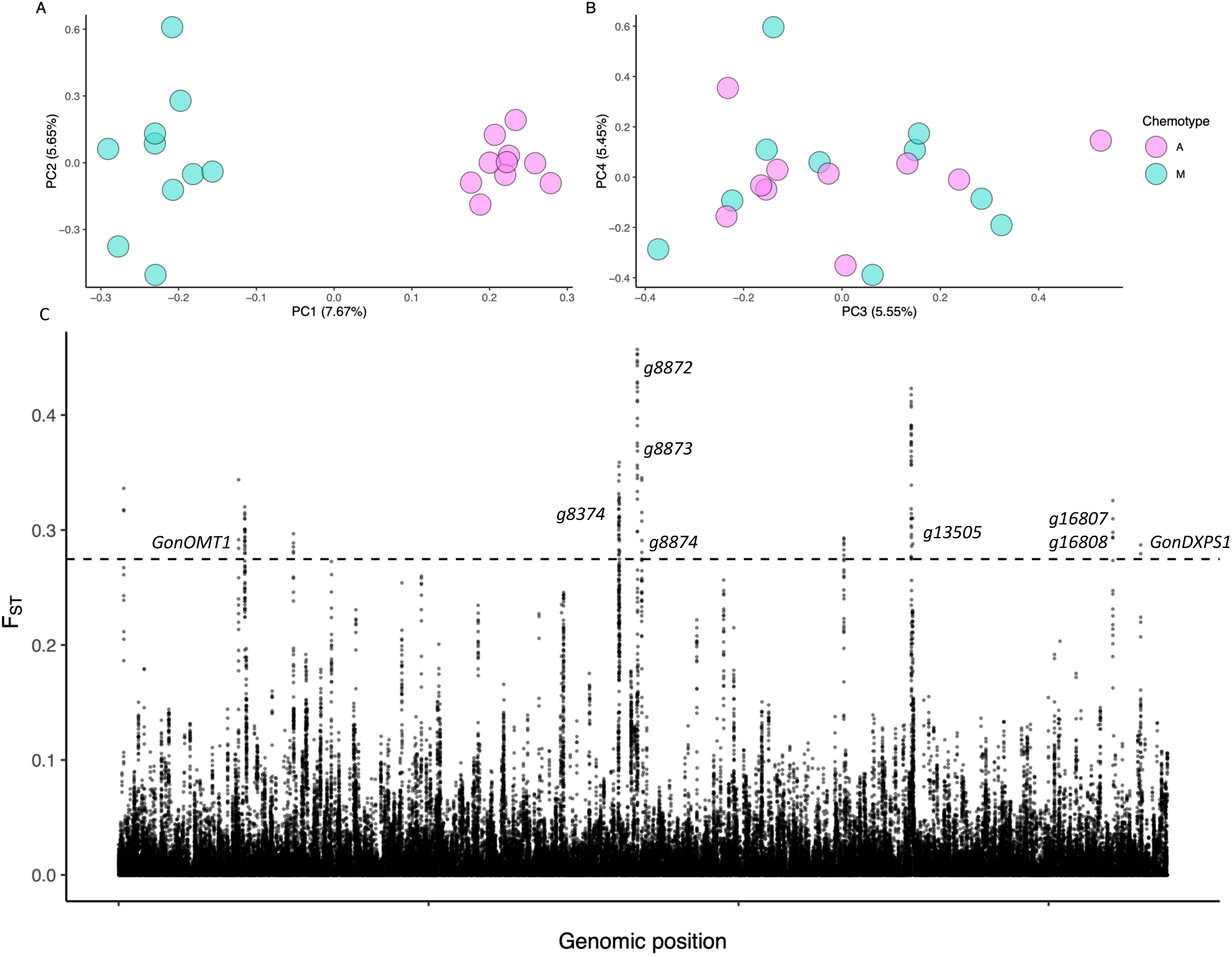
Patterns of genetic variation across the A and M chemotypes. A) and B) show principal components 1 & 2 and 3 & 4, respectively, of genetic variation using unlinked SNPs across the genome. C) Results of a sliding window analysis of F_ST_ between the two chemotypes. The dotted line corresponds to the 99.9th percentile of F_ST_ across these windows (0.275) while labels correspond to annotated genes present within those genomic regions.

Genomic differentiation was concentrated in a relatively small number of regions. Using a sliding window approach, we identified 14 genomic regions containing elevated F_ST_ corresponding to the top 0.1% of values (> 0.275), encompassing nine annotated genes (Fig. 2C, Table 1). Because F_ST_ is prone to erroneously identifying regions as differentiated due to low within-population variance rather than high between-population variance ^23^, we further performed more fine-scale analyses and investigated which of these genes exhibited other signatures of selection (Fig. 3). The region on the scaffold for *g2312*, a putative jasmonic acid carboxyl methyltransferase (*GonOMT1*), showed elevated absolute divergence (D_XY_) in the top 1% of values (> 0.0378; Fig. 3B) in addition to a reduction of heterozygosity in the A chemotype in the top 1% of values (> 0.0125; Fig. 3C). While there were several SNPs near fixation (F_ST_ > 0.9) between the chemotypes around this region, none of them were present within the gene itself (Fig. S2), suggesting divergence in regulatory rather than coding regions. Phylogenetic analysis across plant methyltransferases of diverse functions placed *GonOMT1* in a group of carboxyl methyltransferases closely related to a member responsible for the biosynthesis of floral scent compounds in *Hedychium coronarium* (Fig. 3D, Table S1).

**Figure 3.**
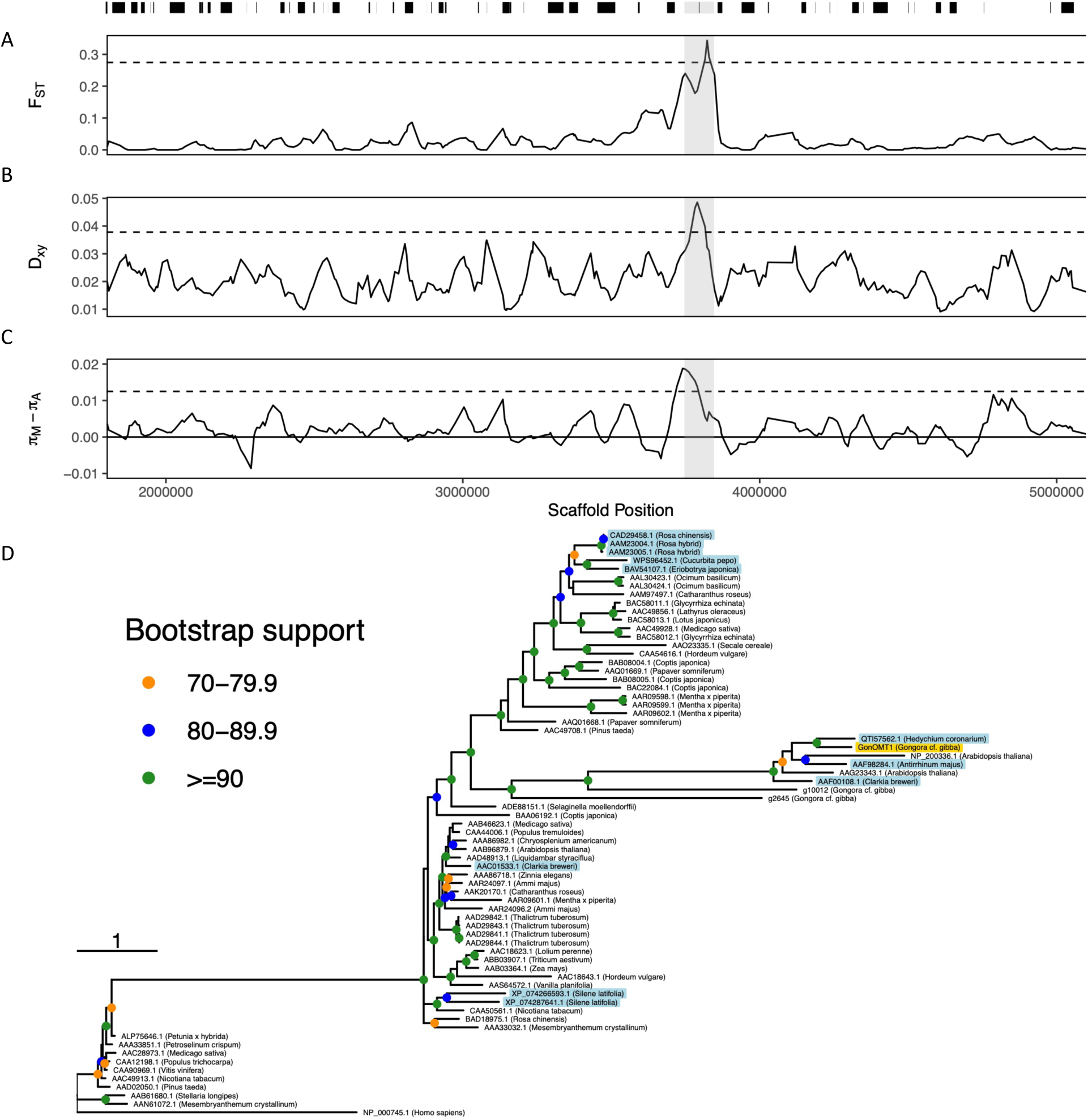
Signatures of selection and evolutionary relationships of *GonOMT1*. A) – C_ show results of sliding windows in the scaffold in which *GonOMT1* is present in. Black bars above show positions of genes in this region, with gray shaded areas corresponding to 50 kb around *GonOMT1*. A) Sliding window estimates of F_ST_, with the dashed line representing the 99.9^th^ percentile of 0.275. B) Sliding window estimates of D_XY_, with the dashed line representing the 99^th^ percentile of 0.0378. C) Sliding window estimates of heterozygosity of A subtracted from heterozygosity of M. The dotted line corresponds to the 99^th^ percentile of this distribution (0.0125). D) A gene tree made from diverse OMTs across plants. Tips highlighted in light blue correspond to OMTs that have been functionally confirmed to play a role in producing floral scent while the tip highlighted in yellow corresponds to *GonOMT1*.

**Table 1.** Genes present in genomic windows identified as extreme F_ST_ outliers (>0.275, the 99.9^th^ percentile)

| Scaffold | Position | Putative function | Annotation name | Gene name in manuscript |
| --- | --- | --- | --- | --- |
| ScYo1bC_2192;HRSCAF=2933 | 3795487 - 3797169 | Jasmonic acid carboxyl methyltransferase | *g2312* | *GonOMT1* |
| ScYo1bC_6133;HRSCAF=8159 | 154256 - 158038 | Pectinesterase | *g8374* |  |
| ScYo1bC_6298;HRSCAF=8377 | 354557 - 354751 | NADH dehydrogenase | *g8872* |  |
| ScYo1bC_6298;HRSCAF=8377 | 554066 - 554297 | Peroxidase | *g8873* |  |
| ScYo1bC_6298;HRSCAF=8377 | 700538 - 701644 | Clathrin assembly protein | *g8874* |  |
| ScYo1bC_7675;HRSCAF=11265 | 668927 - 669226 | Pentatricopeptide repeat-containing protein | *g13505* |  |
| ScYo1bC_8636;HRSCAF=14174 | 172 - 26720 | Boron transporter | *g16806* |  |
| ScYo1bC_8636;HRSCAF=14174 | 28288 - 110333 | YTH domain-containing protein | *g16807* |  |
| ScYo1bC_8830;HRSCAF=14368 | 649970 - 655876 | 1-deoxy-D-xylulose-5-phosphate synthase | *g16995* | *GonDXPS1* |

### Scent-producing floral tissue exhibit upregulation of biosynthetic pathways

As an independent means of characterizing the genetic differences between chemotypes, we performed RNA-sequencing experiment, using floral tissue from 3-5 individuals per chemotype. Uniquely mapping reads after this process ranged from 10.86 million to 25.3 million, with an average of 18.82 million reads per sample (Fig. S1B). In addition to investigating differences between chemotypes, we performed a more fine-scale analysis by comparing two floral tissues – the hypochile, where scent is produced and collected by pollinators, and the epichile which lacks this function (Fig. 1B) ^24,25^. A principal components analysis (PCA) performed across all 26 libraries showed strong clustering by chemotype and tissue, with the first two axes of variation explaining 47% of variation in the data (Fig. 4A).

**Figure 4.**
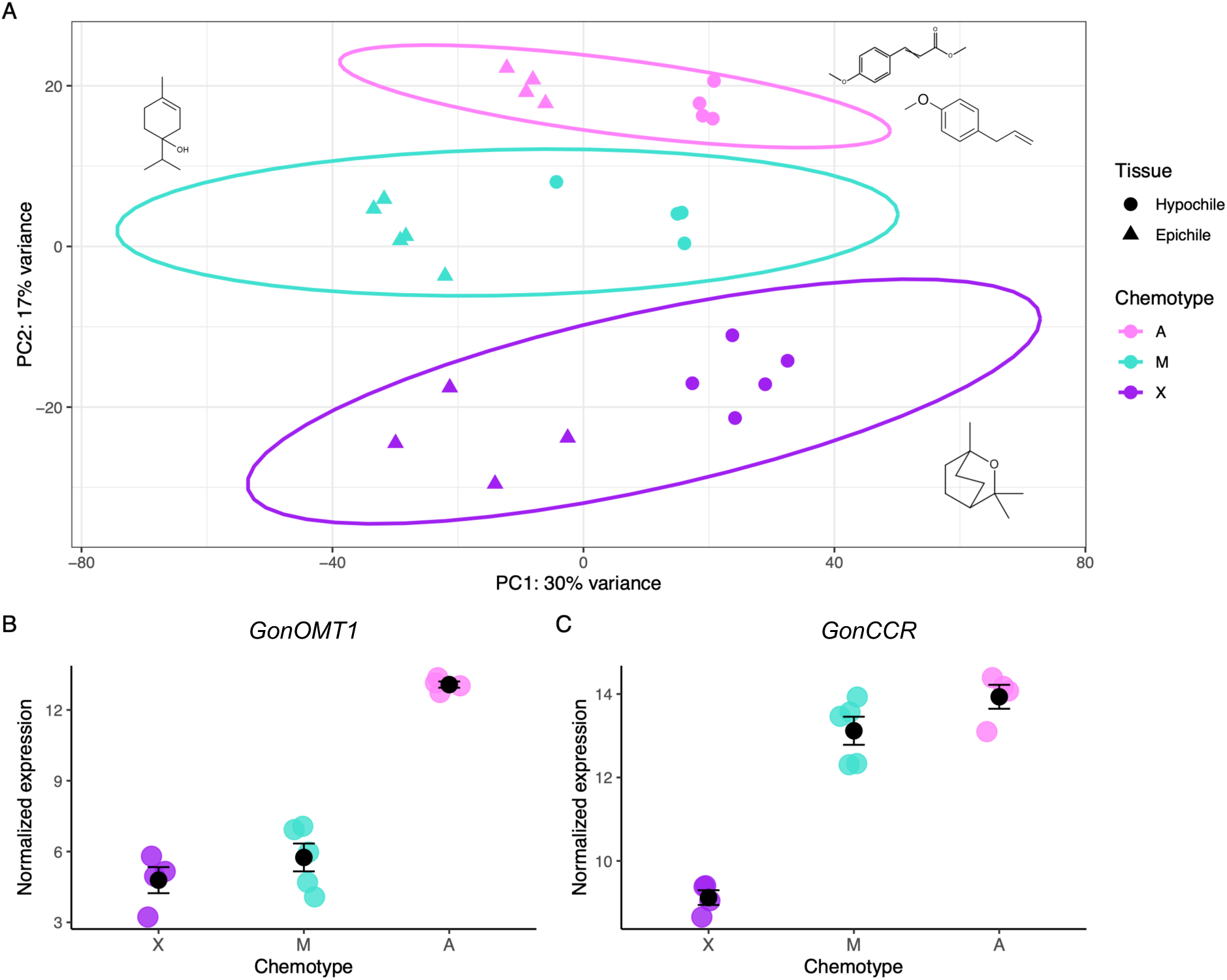
Gene expression patterns across chemotypes. A) PCA across all libraries, with colors corresponding to chemotypes and shapes corresponding to tissue. B) Gene expression patterns in hypochiles across chemotypes of *GonOMT1*. C) Gene expression patterns in hypochiles across chemotypes of *GonCCR*. In the gene expression panels black circles represent means and bars represent standard errors. “Normalized expression” on the y-axes refer to log-transformed expression values following normalization in *DESeq2*.

We identified 2078, 1286, and 689 differentially expressed (DE) genes between the epichile and hypochile tissues within the X, M, and A chemotypes, respectively. Of these genes, 766, 725, and 274, respectively, were upregulated in the hypochile tissue. We found that across these upregulated genes, terms related to terpenoid and phenylpropanoid biosynthesis were significantly enriched (Fig. S3), providing evidence that genes underlying scent production are highly expressed in this tissue. This is consistent with histological studies identifying osmophores – specialized scent-producing cells – in the hypochiles of *Gongora* and related genera of perfume flowers ^24,25^.

### Differentially expressed genes between A and M provide candidates for scent divergence

We identified 1286 DE genes in the A-M contrast. Many terms significantly enriched in the GO analysis were associated with phenylpropanoid synthesis, consistent with our predictions of divergence in biosynthetic pathways at the molecular level (Fig. S3). We found that the most significantly DE gene was *GonOMT1* (Fig. 4B, S3; 892-fold higher in A, P-adj < 0.0001) This gene was further strongly upregulated in the hypochile tissue of the A chemotype (3-fold higher, P-adj < 0.0001).

We found limited evidence of strong downregulation of TPS genes in the A chemotype, with no significant enrichment of terms associated with terpene synthesis in the upregulated genes from M in this chemotype. However, we found that *g16995*, present in a genomic region of strong differentiation, was also significantly upregulated in the M chemotype relative to the A chemotype (3.1-fold higher, P-adj < 0.0001). This gene is a putative 1-deoxy-d-xylulose-5- phosphate synthase (*GonDXPS1*), coding for an enzyme responsible for the initial rate-limiting step of the MEP pathway, a highly conserved pathway for synthesis of isoprenoid precursors of monoterpenes ^26,27^. In addition to strong differential expression between the A and M chemotypes, this gene exhibited tissue-specific expression, with significant upregulation in the hypochiles of all 3 chemotypes. The only significantly differentially expressed TPS from this contrast was *GonTPS3*, which was upregulated in A (33.6-fold higher, P-adj = 1.6 x 10^-4^). This TPS was previously annotated as a member of the “TPS-a” clade, a group of TPSs typically responsible for sesquiterpene production, consistent with the presence of these compounds as a minor component of A’s volatile blend^16,28^.

### Differentially expressed genes between M and X provide insight into divergence across two biosynthetic pathways

We identified 1166 DE genes in the M-X contrast. We found substantial enrichment of GO terms associated with terpene synthesis, including several for specific monoterpenes (Fig. S3). As TPS’s catalyze the final step of this biosynthetic pathway, determining compound identity ^18^, we expected that they would be strongly DE in this contrast comparing two chemotypes with differing monoterpene scent profiles. We indeed found that *GonTPS18*, a putative monoterpene synthase previously annotated as a member of the “e/f-clade” ^28^, was substantially differentially expressed, with over a thousand-fold higher expression (P-adj = 4.03 x 10^-4^) in the X chemotype. Additionally, we found that several terms corresponding to terpene synthesis including “dimethylallyl diphosphate metabolic process” and “R-linalool synthase activity” were significantly enriched in upregulated genes from the X chemotype (P = 0.028 and 0.04, respectively).

One of the most strongly enriched terms in the upregulated genes from the M chemotype was “phenylpropanoid biosynthetic process” (P = 0.0041; Fig. S3), suggesting strong differences in expression of this metabolic pathway despite not being represented in the phenotypic differences. Furthermore, the most significantly upregulated gene in the M chemotype was *g12713*, a putative cinnamoyl co-A reductase (Fig. 4C; *GonCCR*; 63.9-fold higher in M, P-adj < 0.0001). Enzymes encoded by these genes catalyze a critical step in phenylpropanoid biosynthesis, directing metabolic flux towards the production of monolignols ^20,29^.

### Co-expression networks reveal coordinated transcriptional programs across chemotypes and tissues

To provide further context for the candidate genes identified in the population genetic and transcriptomic analyses, we performed weighted gene correlation network analysis (WGCNA) that could potentially reveal clusters of co-expressed genes (“modules”) underlying biosynthetic divergence (Fig. S4). We specifically searched for modules that the candidate genes were members of and identified highly connected members of these modules (“hub genes”) that might correspond to factors regulating major shifts in transcriptional dynamics underlying transitions in floral scent phenotypes.

*GonOMT1* was present in a module of 454 genes with high expression specific to the A chemotype (Fig. 5, henceforth referred to as the “aromatic modification module”). This module contained 16.7% of the upregulated genes in A from the A-M contast with over 6-fold more genes from this category than expected by chance (Fig. S5) and was enriched for GO terms related to aspects of phenylpropanoid synthesis, such as “flavonoid biosynthetic process” (P = 0.00205) and “L-phenylalanine catabolic process” (P = 0.00522). *GonOMT1* was present as a hub gene in the module, suggesting its coordinated expression in the core pathway of the network. Another hub gene present in this module was *g14202*, a putative basic helix-loop-helix (bHLH) transcription factor. This gene exhibited no significant differential expression between the M and X chemotypes but was upregulated 4.3-fold in the A chemotype compared to the M chemotype (P-adj < 0.0001).

**Figure 5.**
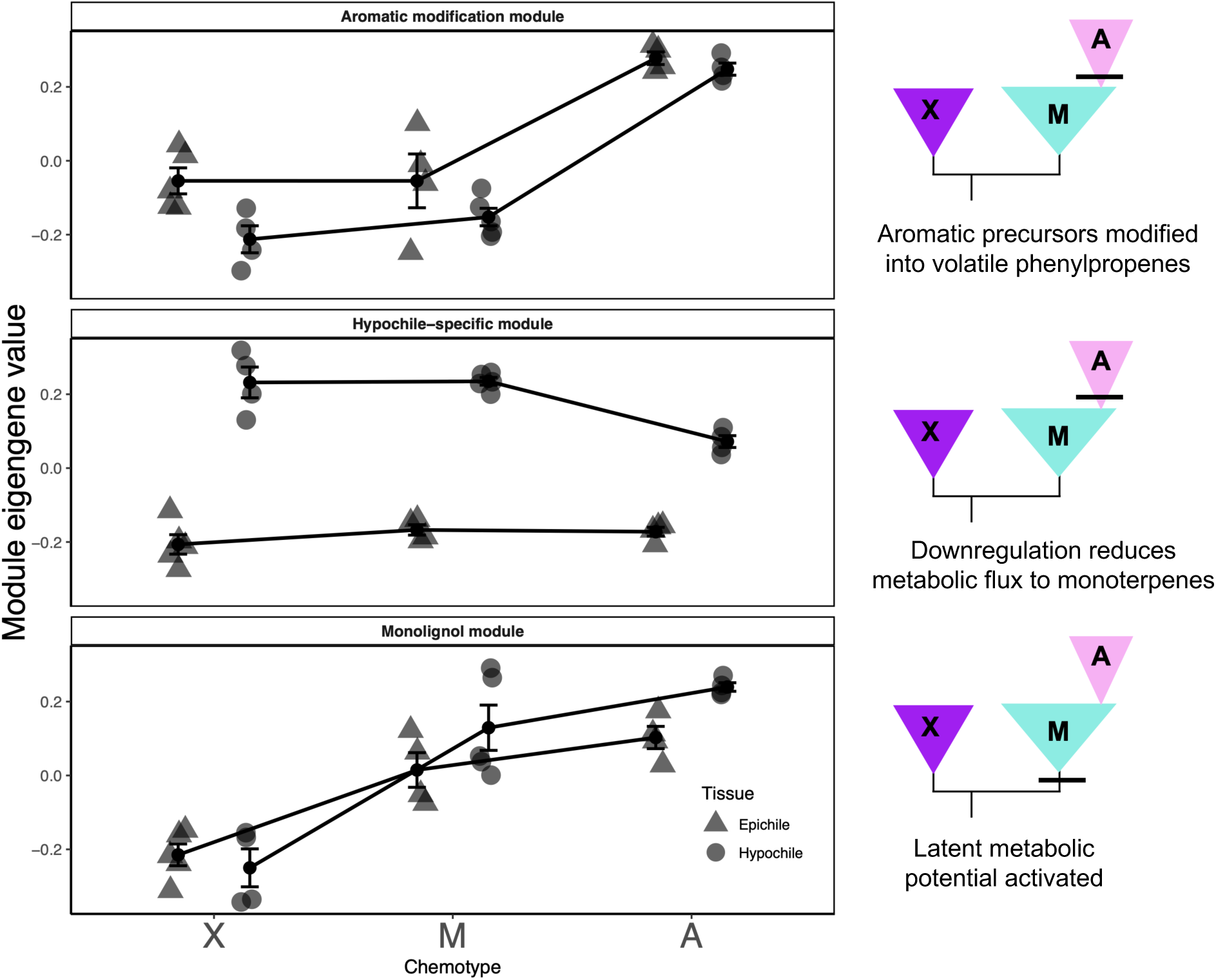
Conceptual model of how coordinated patterns of gene expression shape scent evolution via contingent processes – with activation of the monolignol module in the ancestor of the A-M clade, followed by novel modification in the hypochile-specific and aromatic modification modules in the A lineage.

*GonDXPS1* was present in a larger module of 1380 genes exhibiting weak association with chemotypes but a strong association with tissue, with much higher expression in the hypochile (henceforth referred to as the “Hypochile-specific module”). This module exhibited strong enrichment of genes involved in biosynthetic pathways relevant to scent production such as “chorismate biosynthetic process” (P < 0.0001) and “terpenoid biosynthetic process” (P = 0.0001), suggesting a broad importance in regulating floral secondary metabolites. *GonDXPS1* was present as a hub gene, as was a putative MYB20 transcription factor *g978*. MYB20s are related to *ODORANT1*, a master regulator of floral volatile production in *Petunia*, providing a potential candidate for a driver of this altered transcriptional pattern ^30,31^. This gene exhibited 5.2-fold downregulation in the A chemotype compared to the M chemotype (P-adj < 0.0001), with significant upregulation in the hypochiles within each chemotype (2.8 – 12.1-fold upregulation, all P-adj < 0.0001).

*GonCCR* was present in a module of 380 genes exhibiting a pattern of low expression in the X chemotype, moderate levels in the M chemotype, and highest expression in the A chemotype (Fig. 5, S4,6; henceforth referred to as the “monolignol module”). This module contained 11.7% of the upregulated genes in M from the M-X contrast, with over 5-fold more than expected by chance (Fig. S5), and was enriched for processes related to monolignol biosynthesis such as “lignin biosynthetic process” (P = 2.4 x 10^-4^) and “phenylpropanoid metabolic process” (P = 0.02057). *GonCCR* was present as a hub gene, along with a putative TCP14 transcription factor *g1942*. This gene was expressed 2.19-fold higher in A compared to M (P-adj < 0.0001) and 8.2-fold higher in M compared to X (P-adj = 5.8 x 10^-4^).

## Discussion

Floral scent is a highly evolutionarily labile signaling trait that can contribute to reproductive isolation in plants, often evolving rapidly in the process of speciation. Transitions between biosynthetic pathways are a common mode of floral scent divergence and provide a lens in which to understand how evolutionary processes underlie major patterns of phenotypic diversity. By performing population genomic and transcriptomic analyses across a rapid transition coupled with gene expression analysis across a deeper divergence (Fig. 1A), we provide evidence that contingency may facilitate novel suites of gene expression, allowing disproportionately large phenotypic changes to arise from limited genomic divergence (Fig. 5).

Production of the novel aromatic-dominated phenotype in the A chemotype from a terpene-emitting background is the result of two divergent processes – recruitment of aromatic biosynthesis coupled with the loss of terpene production. *GonOMT1* emerged as a strong candidate contributing to the former process. This gene was present in a locus exhibiting strong signatures of selection, while further exhibiting both tissue-specific and high expression in the A chemotype compared to the M chemotype. OMTs are a large family of enzymes that methylate oxygen atoms in secondary metabolites to play key roles in a variety of processes in plants ^32^, while those involved in floral volatile evolution often modify the final step of biosynthesis to generate divergent products ^33,34^. *GonOMT1* was phylogenetically related to salicylic acid carboxyl methyltransferases that have confirmed function in floral scent production in other plant species (Fig. 3D) ^35,36^. While the recruitment of OMTs to floral scent production has been found to involve positive selection on coding regions following gene duplication ^37^, we found no fixed SNPs in *GonOMT1* between chemotypes. In addition, given the drastic 892-fold upregulation in A, our results instead suggest regulatory divergence as a potential mechanism underlying *GonOMT1* recruitment, although future functional experiments are required to more rigorously test this hypothesis.

*GonOMT1* was present within a module of co-expressed genes differentiating the terpene-producing chemotypes from the A chemotype. This gene was a hub within this module, indicating patterns of expression highly correlated with a broader transcriptional shift distinguishing A from the monoterpene-emitting chemotypes. The presence of *g14202*, a putative bHLH transcription factor, among these hub genes reveals a potential regulatory candidate that could coordinate the expression of this network as they have been found to regulate floral scent production in other species ^38,39^. While population genetic and RNA-sequencing allowed us to identify these candidate genes and networks potentially underlying scent divergence between A and M, these changes alone are likely insufficient to generate scent divergence due to the major differences in biosynthetic pathways and OMTs tending to function towards the end of floral scent biosynthesis. We hypothesized that there may have been upregulation of biosynthetic machinery in M that predisposed this lineage towards a shift in floral scent.

Despite both producing floral scent profiles dominated by cyclic monoterpenes with no aromatic compounds, we identified upregulation of phenylpropanoid pathway genes in the M chemotype compared to the X chemotype, with *GonCCR* being the most upregulated gene in M from this contrast. CCRs perform the first committed step in monolignol biosynthesis, directing metabolic flux towards this pathway^29^. Expression of this gene in the lineage leading to the A-M group could have occurred via evolutionary processes unrelated to floral scent production, such as in structural reinforcement of tissue or production of defense compounds. The resultant alterations of metabolic flux could therefore provide substrates for downstream phenylpropanoid enzymes that were subsequently recruited into volatile aromatic production in the A chemotype (Fig. 5). Network analysis provided complementary support for this interpretation, with *GonCCR* present as a hub gene in the “monolignol module” exhibiting low expression in X, upregulation in M, and substantial upregulation in A. Many of the other hub genes from this module were components of the phenylpropanoid pathway and exhibited upregulation in the M chemotype relative to the X, suggesting a concerted shift towards this biosynthesis along this lineage. Together, these findings support a model of evolutionary contingency, where pre-existing transcriptional activity facilitated divergence following upregulation of a key enzyme. Such a mechanism may explain the prolific shifts between terpene- and aromatic-dominated profiles observed across perfume flower lineages^11^, highlighting how latent metabolic potential can shape evolutionary trajectories.

Downregulation of *GonDXPS1*, the first enzyme in the MEP pathway, may have contributed to the loss of monoterpenes in A by reducing the production of their isoprenoid precursors. Experimental silencing of DXPS has been found to modify tissue-specific patterns of volatile monoterpene production ^43,45^. Notably, increased production of sesquiterpene compounds was noted after knocking out a DXPS in *Solanum lycopersicum*, likely due to a shift in metabolic flux from the MEP pathway to the mevalonate pathway, a parallel pathway responsible for sesquiterpene precursors. This mirrors the chemical phenotypes of the A vs. M chemotypes, where the A chemotype lacks monoterpenes but produces a small quantity of sesquiterpenes. While *GonDXPS1* did not exhibit signatures of strong selection, it was located in a genomic region of elevated F_ST_ and substantially higher diversity in A compared to M, potentially a result of relaxed selective constraint on this gene following reduced functional importance of this gene (Fig. S7). This gene was a hub in a hypochile-specific module strongly enriched for biosynthetic activity that was additionally downregulated in the hypochiles of A, providing further evidence for its role in shaping scent phenotypes.

While changes in expression levels of TPSs and divergence in their coding sequences have been identified as drivers of floral scent evolution and pollinator shifts between flowering plant species, including in orchids ^40,41^, we identified mixed evidence for their role in shaping scent divergence across *Gongora* chemotypes. A putative monoterpene synthase with substantial in the X chemotype could be responsible for the divergence in monoterpene composition between M and X. This TPS, *GonTPS18*, may provide insight into the production of 1,8-cineole, the dominant compound in X and the most widespread compound across perfume flowers^11^. Experimental validation of this candidate gene and TPSs expressed across other perfume flowers will provide insight into the repeated evolution of this compound across disparate lineages. By contrast, we found limited evidence of TPS downregulation or modification driving loss of monoterpene emission between the A and M chemotypes, with a lack of strong signal from both the population genetic and transcriptomic analyses. These results contrast with other systems where downregulation^42^ and pseudogenization^44^ of TPSs were found in concordance with loss of floral scent emission.

We suggest several lines of future work at different scales of biological organization that would provide further insight into our findings. At the molecular level, functional characterization of candidate enzymes we identified, especially *GonOMT1*, *GonCCR*, *GonTPS18*, and *GonTPS3*, will provide more direct tests of how they are involved in biosynthetic processes. Our co-expression analyses also identified several transcription factors as candidate regulators of metabolic programs involved in scent, including members of bHLH, MYB, and TCP families. Characterization of the transcription factors we identified via DNA-binding assays will be valuable for confirming their role in activating the genes of interest. Another natural follow-up to this research would be to perform whole genome sequencing of the X chemotype. Providing such an outgroup would allow for greater understanding of the dynamics of introgression across these *Gongora* populations and provide potential genomic insights into the demographic processes leading to labile chemical evolution in the A-M group. Finally, at the macroevolutionary scale, a concerted effort to sample across independent transitions between aromatic- and terpenoid-dominated species or chemotypes would provide clearer insight into the pervasiveness of contingency as a mechanism for increasing the speed at which the transitions occur.

Together, our results suggest that the evolutionary accessibility of a novel floral scent phenotype can depend on the molecular state inherited from its ancestor. By sampling across several reproductively-isolated chemotypes of *Gongora*, we provide a more comprehensive view of how this sexual signaling trait evolves so rapidly. Elevated expression of phenylpropanoid biosynthetic genes in the monoterpene-emitting M chemotype preceded the emergence of the aromatic-emitting A chemotype, while subsequent changes in pathway regulation accompanied rapid divergence in floral volatiles and pollinator associations. These patterns support evolutionary contingency as a potential mechanism by which disproportionately large phenotype differences, and consequently, reproductive isolation, can arise via relatively few genetic changes. More broadly, our findings suggest that latent metabolic potential can affect the adaptive landscape, facilitating certain phenotypic transitions.

## Materials and Methods

### Scent Extraction

In addition to a first pass of identification by smelling flowers, we used previously established methods to extract floral scent from these chemotypes (see Hetherington-Rauth and Ramírez 2016 and Guizar Amador 2022 for full details). Briefly, inflorescences were enclosed in an oven bag for 30 minutes, then air was pumped out at 500 mL/hour through 2 cm long glass capillary tubes filled with Tenax and charcoal (“scent traps”) for an hour. Scent traps were then eluted using 200 uL of hexane into Agilent vials with glass inserts and stored at −20 degrees C until analysis. We used an Agilent 7890B GC fitted with a 30m x 0.25mm x 0.25mm HP-5 Ultra inert column, coupled to an Agilent 5977A mass spectrometer (Agilent Technologies). Relative proportions of compounds (%) were calculated by dividing the absolute amounts of individual compounds by the sum of all compounds.

### DNA Extraction

Fresh leaf tissue of 10 plants each of the A and M chemotypes were collected from previously collected plants growing at La Gamba Biological Station in Puntarenas Province, Costa Rica (Guizar Amador 2022). Genomic DNA extraction was performed using the Plant DNEasy Mini Kit (Qiagen, Germany) and extractions were sent to Novogene for library construction and whole genome resequencing on the NovaSeq 6000 platform to generate 150 bp paired end reads to a target coverage of 8x.

### Population Genetics

After assessing the quality of the sequencing data with FASTQC v.0.11.7, the reads were mapped to the *Gongora* cf. *gibba* (chemotype A) reference genome using BWA-MEM, as implemented in BWA v.0.7.18 ^28,46^. Samtools v.1.17 was then used to mark duplicates and index the bam files ^47^. We used bcftools v.1.16 to call variants and filtered them with vcftools v.1.16 to remove insertions and deletions, sites with minor allele frequency under 0.08, sites with over 5% missing data, sites with mapping quality below 30, and sites with a sequencing depth below 5x or over 20x ^48^.

To investigate population structure between the chemotypes, we used plink v1.07 to perform a PCA based on 290,366 unlinked sites ^49^. To estimate genome wide F_ST_, we used vcftools to compute Weir F_ST_ across all 7,124,345 filtered SNPs. To understand broader patterns of divergence and nucleotide diversity, we used the popgenWindows.py script written by Simon Martin (https://github.com/simonhmartin/genomics_general/blob/master/popgenWindows.py) to estimate F_ST_, D_XY_ and pi over 50kb sliding windows with a 10kb step size and a minimum of 1kb sites covered.

### Gene Tree Analyses

We compiled sequences of OMTs from a previous study investigating gene family evolution of the group, in addition to several additional functionally characterized OMTs involved in floral scent production (Table S1; Pak et al. 2004; Lam et al. 2007; Gupta et al. 2012; Koeduka et al. 2016; Yue et al. 2021; Hoepflinger et al. 2024) and a human OMT as an outgroup. We aligned the sequences using MAFFT v.7.562^52^ and then used IQ-TREE 2^53^ to infer the phylogenetic relationships, using default settings for proteins with 1000 bootstraps. We plotted this gene tree using ggtree^54^.

### RNA extraction

We collected floral tissue from the previously-described plants between March and May of 2022 and 2023. Fresh floral tissue on the first day of anthesis was collected between 9 and 9:30 AM local time and was dissected into the hypochile and epichile, with the former region corresponding to the source of scent production and collection by bees ^25^. Tissue samples were placed into 1.5 mL microcentrifuge tubes with sterile stainless steel beads and immediately flash frozen in liquid nitrogen until extraction.

RNA from tissue was extracted using Qiagen RNEasy Mini kits (Qiagen, German), with frozen floral samples first disrupted with a TissueLyser and homogenized in Qiashredders. Total RNA was eluted into 30 microliters of reagent grade water before dried in Gentegra tubes for storage. When samples were ready for sequencing, the same volume of reagent grade water was used to rehydrate the samples before they were sent to Novogene for quality control, library preparation, and sequencing on a NovaSeq 6000 platform using 150 bp paired-end reads. A total of 26 samples were used in the analyses presented, with 9 from the X chemotype, 9 from the M chemotype, and 8 from the A chemotype.

### Data processing

We used TrimGalore!^55^ to filter out low quality sequences with Phred scores of below 20 and remove adaptors. We then ran 2-pass mapping of retained reads to the *Gongora* cf. *gibba* (chemotype A) reference genome (Guizar Amador et al. 2024) using STAR v 2.7.1^56^. We then counted mapped reads using featureCounts ^57^ to generate a text file output read into R v.4.2.1 for downstream analyses. We performed principal components analysis (PCA) on a variance-stabilized transform of the dataset (a method to normalize expression of transcripts across a heterogeneous sample) to visualize the data prior to differential expression analyses and identify clear patterns of clustering.

### Differential expression analysis between chemotypes

We 2 categories of contrasts – comparisons within chemotypes between the two tissues and comparisons of hypochile expression between chemotypes. The former set of comparisons were performed to confirm increased expression of genes related to scent production in the hypochiles while the latter set were performed to capture genes that may underlie divergence in scent production between populations. Specifically, the M-X contrast captures divergence within the terpene synthesis pathway, contrasting two more deeply diverged populations varying with respect to the monoterpenes they produce. The A-M contrast captures divergence between biosynthetic pathways and the primary axis of variation across perfume flowers, comparing populations that are very shallowly diverged.

To compare differentially expressed (DE) genes within each contrast, we used DESeq2 v.1.36 ^58^. We first filtered out genes with fewer than 10 transcripts in any sample to reduce noise from low-abundance transcripts. We defined significantly DE genes as those with an absolute fold change of at least 2 and a P-value adjusted for multiple comparisons (P-adj) below 0.05.

To characterize pathways that may have been enriched in these comparisons, we performed gene ontology (GO) analysis to compare annotations of differential genes against background expression. We used BLASTP of the Gongora proteome against the swiss-prot database implemented in DIAMOND v. 2.0.10, then combined this annotations with additional annotations from InterPro to improve the percentage of genes with GO annotations in our analysis ^59,60^. We used BLAST2GO map the annotations and imported these functional annotations into R to carry out enrichment analysis with topGO ^61^. We considered the “background” to be all expressed genes considered in the specific differential expression analysis after filtering.

### Gene network analysis

While DE analyses provide a starting point towards identifying genes functionally relevant to a scent divergence based on changes in expression levels, a strong limitation for DE analysis is that each gene is considered in isolation while genes more realistically function in networks. For example, switching between biosynthetic pathways involved in scent production likely involves major changes in regulatory architecture where suites of genes - “genetic modules” - may shift their patterns of co-expression, making standard differential expression analysis of individual genes alone insufficient to understanding overall phenotypic divergence. To discover these genetic modules associated with scent differentiation, we conducted a weighted gene co-expression network analysis (WGCNA) across our transcriptomic samples ^62^.

We used the R package WGCNA v1.72 to identify co-expressed modules of genes across all samples. We first normalized expression of genes across samples using the varianceStabilizingTransformation function from DESeq2. The adjacency matrix was then calculated using a soft thresholding power of 11 based on the scale-free topology fit index (R^2^ = 0.8) calculated using the pickSoftThreshold function. In contrast to a hard threshold, of which correlation values above or below a certain value are considered connected or not, respectively, the soft threshold raises the correlation to a power so that differences between strong and weak correlations are exaggerated rather than defined into binary terms. The topological overlap matrix was calculated using a deepSplit value of 2, minModuleSize was set to 30, and mergeCutHeight was set to 0.25. To investigate the association between the modules with phenotypes, we computed their “eigengene” values (i.e. a value analogous to the first principal component of expression levels within a module, providing an abstraction of overall expression patterns) and correlated them with chemotypes.

To identify potential “hub genes” that may be coordinating these larger patterns of co-expression, we selected genes that both exhibited the top 20% of connectivity within their modules and had absolute values of module membership (i.e. exhibiting high correlation with overall module expression) above 0.75.

## Acknowledgments

The authors would like to extend our gratitude to the staff at La Gamba Biological Station for facilitating the field work. Additionally, the authors would like to thank K. M. Kay, C. P. Gross, and members of the Ramírez and Espíndola Labs (M.C. Sandoval, P. Coggan, D.Y. Ramírez, J. El-Hifnawi, A. Espíndola, J. O’Neill) for helpful comments on early versions of the manuscript. The Comisión Institucional de Biodiversidad of the University of Costa Rica (CBio-UCR) issued the resolution authorizing access to genetic and biochemical resources under the project “C4074 Phylogenomics, morphometrics, and taxonomy of the genus *Gongora* (Orchidaceae) in Costa Rica.”

## Notes

### Competing Interest Statement

The authors have declared no competing interest.

### Funding

- NSF
- Davis Botanical Society
- Center for Population Biology
- David and Lucile Packard Foundation

