## Supplemental Materials for "Evolutionary contingency facilitates scent divergence in an emergent orchid population"

**Table S1.** O-methyltransferases used in phylogenetic analysis

| Accession | Name | Species | Floral volatile |
|---|---|---|---|
| AAQ01669.1 | (RS)-norcoclaurine 6-O-methyltransferase | Papaver somniferum | |
| AAQ01668.1 | (R,S)-reticuline 7-O-methyltransferase | Papaver somniferum | |
| BAA06192.1 | S-adenosyl-L-methionine:scoulerine 9-O-methyltransferase | Coptis japonica | |
| BAB08005.1 | S-adenosyl-L-methionine:3'-hydroxy-N-methylcoclaurine 4'-O-methyltransferase | Coptis japonica | |
| BAC22084.1 | columbamine O-methyltransferase | Coptis japonica | |
| BAB08004.1 | S-adenosyl-L-methionine:norcoclaurine 6-O-methyltransferase | Coptis japonica | |
| ABB03907.1 | flavonoid O-methyltransferase | Triticum aestivum | |
| AAM97497.1 | flavonoid O-methyltransferase | Catharanthus roseus | |
| CAA54616.1 | flavonoid 7-O-methyltransferase | Hordeum vulgare | |
| AAC49856.1 | 6a-hydroxymaackiain methyltransferase | Lathyrus oleraceus | |
| BAC58012.1 | S-adenosyl-L-methionine: daidzein 7-0-methyltransferase | Glycyrrhiza echinata | |
| BAC58013.1 | S-adenosyl-L-methionine: 2,7,4'-trihydroxyisoflavanone 4'-O-methyltransferase | Lotus japonicus | |
| BAC58011.1 | S-adenosyl-L-methionine: 2,7,4'-trihydroxyisoflavanone 4'-O-methyltransferase | Glycyrrhiza echinata | |
| AAC49928.1 | isoflavone-O-methytransferase | Medicago sativa | |
| AAR09602.1 | flavonoid 4'-O-methyltransferase | Mentha x piperita | |
| AAR09601.1 | flavonoid 3'-O-methyltransferase | Mentha x piperita | |
| AAR09599.1 | flavonoid 7-O-methyltransferase | Mentha x piperita | |
| AAR09598.1 | flavonoid 7-O-methyltransferase | Mentha x piperita | |
| AAB96879.1 | O-methyltransferase 1 | Arabidopsis thaliana | |
| AAA33032.1 | myo-inositol O-methyl transferase | Mesembryanthemum crystallinum | |
| AAF00108.1 | S-adenosyl-L-methionine:salicylic acid carboxyl methyltransferase | Clarkia breweri | yes |
| AAF98284.1 | SAM:benzoic acid carboxyl methyltransferase | Antirrhinum majus | yes |
| AAG23343.1 | S-adenosyl-L-methionine:jasmonic acid carboxyl methyltransferase | Arabidopsis thaliana | |
| NP_200336.1 | IAA carboxylmethyltransferase 1 | Arabidopsis thaliana | |
| AAM23005.1 | orcinol O-methyltransferase | Rosa hybrid | yes |
| AAM23004.1 | orcinol O-methyltransferase | Rosa hybrid | yes |
| BAD18975.1 | phloroglucinol O-methyltransferase | Rosa chinensis | |
| CAA50561.1 | catechol O-methyltransferase | Nicotiana tabacum | |
| AAK20170.1 | caffeic acid O-methyltransferase | Catharanthus roseus | |
| AAR24096.2 | bergaptol O-methyltransferase | Ammi majus | |
| AAR24097.1 | caffeic acid O-methyltransferase | Ammi majus | |
| AAL30424.1 | eugenol O-methyltransferase | Ocimum basilicum | |
| AAL30423.1 | chavicol O-methyltransferase | Ocimum basilicum | |
| AAD48913.1 | caffeate O-methyltransferase | Liquidambar styraciflua | |
| AAO23335.1 | O-methyltransferase | Secale cereale | |
| CAA44006.1 | lignin bispecific acid/5-hydroxyferulic acid methyltransferase | Populus tremuloides | |
| AAB03364.1 | O-methyltransferase | Zea mays | |
| AAC18623.1 | bispecific caffeic acid/5-hydroxyferulic acid O-methyltransferase | Lolium perenne | |
| AAC01533.1 | SAM:(Iso)eugenol O-methyltransferase | Clarkia breweri | |
| AAD29844.1 | catechol O-methyltransferase | Thalictrum tuberosum | |
| AAD29843.1 | catechol O-methyltransferase | Thalictrum tuberosum | |
| AAD29842.1 | catechol O-methyltransferase | Thalictrum tuberosum | |
| AAD29841.1 | catechol O-methyltransferase | Thalictrum tuberosum | |
| AAB46623.1 | S-adenosyl-L-methionine: caffeic acid 3-0-methyltransferase | Medicago sativa | |
| AAC49708.1 | caffeic acid O-methyltransferase | Pinus taeda | |
| AAC18643.1 | N-methyltransferase | Hordeum vulgare | |
| AAA86718.1 | S-adenosyl-L-methionine:caffeic acid 3-O-methyltransferase | Zinnia elegans | |
| AAA86982.1 | caffeic acid O-methyl transferase | Chryosplenium americanum | |
| AAN61072.1 | O-methyltransferase | Mesembryanthemum crystallinum | |
| AAB61680.1 | S-adenosyl-L-methionine:trans-caffeoyl-CoA 3-O-methyltransferase | Stellaria longipes | |
| AAC49913.1 | caffeoyl-coenzymeA O-methyltransferase | Nicotiana tabacum | |
| AAD02050.1 | caffeoyl-CoA O-methyltransferase | Pinus taeda | |
| AAA33851.1 | caffeoyl-CoA 3-O-methyltransferase | Petroselinum crispum | |
| CAA12198.1 | caffeoyl-CoA 3-O-methyltransferase | Populus trichocarpa | |
| CAA90969.1 | caffeoyl-CoA O-methyltransferase | Vitis vinifera | |
| AAC28973.1 | S-adenosyl-L-methionine:trans-caffeoyl-CoA 3-O-methyltransferase | Medicago sativa | |
| g2312.t1CDS=1-1113 | GonOMT1 | Gongora cf. gibba | |
| g2645.t1CDS=1-1272 | g2645 | Gongora cf. gibba | |
| g10012.t1CDS=1-1740 | g10012 | Gongora cf. gibba | |
| ALP75646.1 | caffeoyl CoA O-methyltransferase | Petunia x hybrida | |
| ADE88151.1 | caffeyl alcohol/5-hydroxyconiferyl alcohol 3/5-O-methyltransferase | Selaginella moellendorffii | |
| CAD29458.1 | orcinol O-methyltransferase | Rosa chinensis | yes |
| AAS64572.1 | caffeic acid O-methyltransferase | Vanilla planifolia | |
| XP_074287641.1 | cathecol O-methyltransferase 1-like | Silene latifolia | yes |
| XP_074266593.1 | cathecol O-methyltransferase 1-like | Silene latifolia | yes |
| WPS96452.1 | 4-methoxyphenol-O-methyltransferase | Cucurbita pepo | yes |
| QTI57562.1 | benzoic acid/salicylic acid carboxyl methyltransferase 2 | Hedychium coronarium | yes |
| NP_000745.1 | catechol O-methyltransferase | Homo sapiens | |

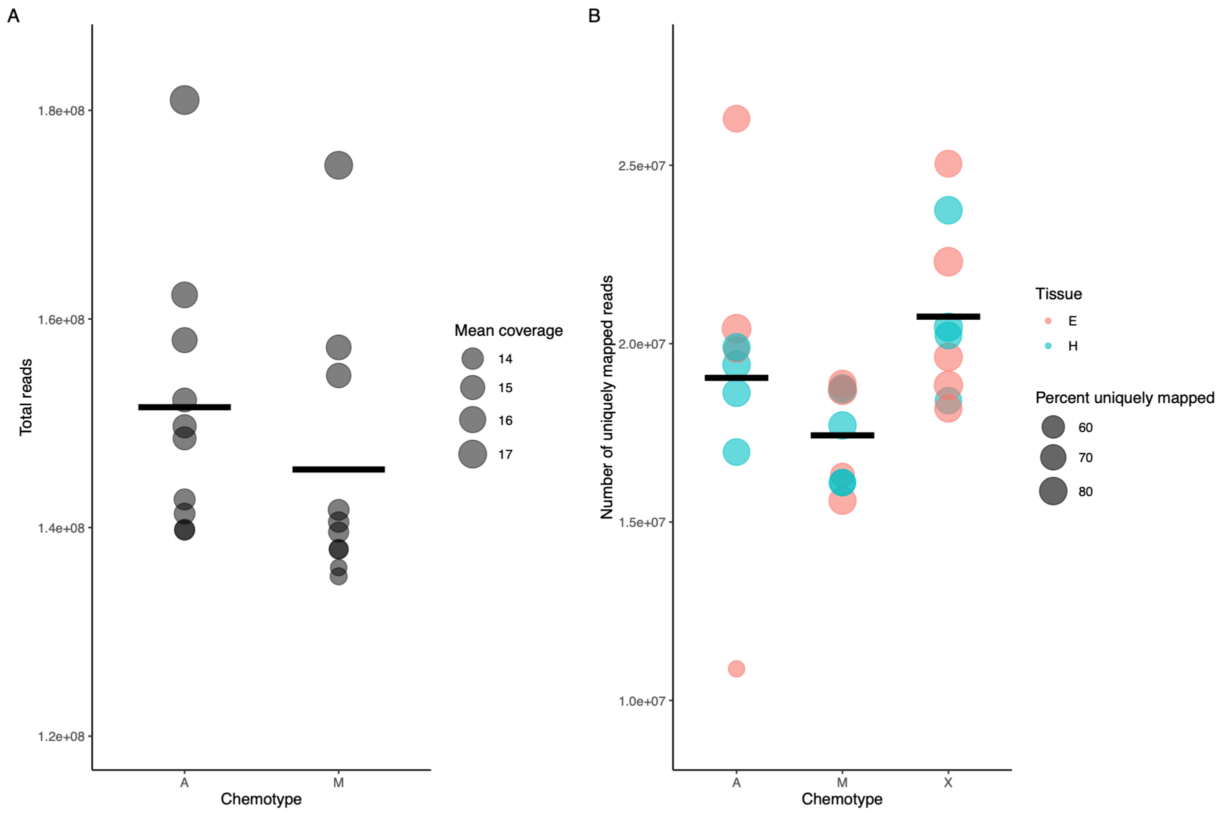


**Fig. S1.** Sample info for A) Whole genome sequencing data and B) RNA-sequencing data, where each point corresponds to a different library. All differences between chemotypes are non-significant ($P > 0.05$).

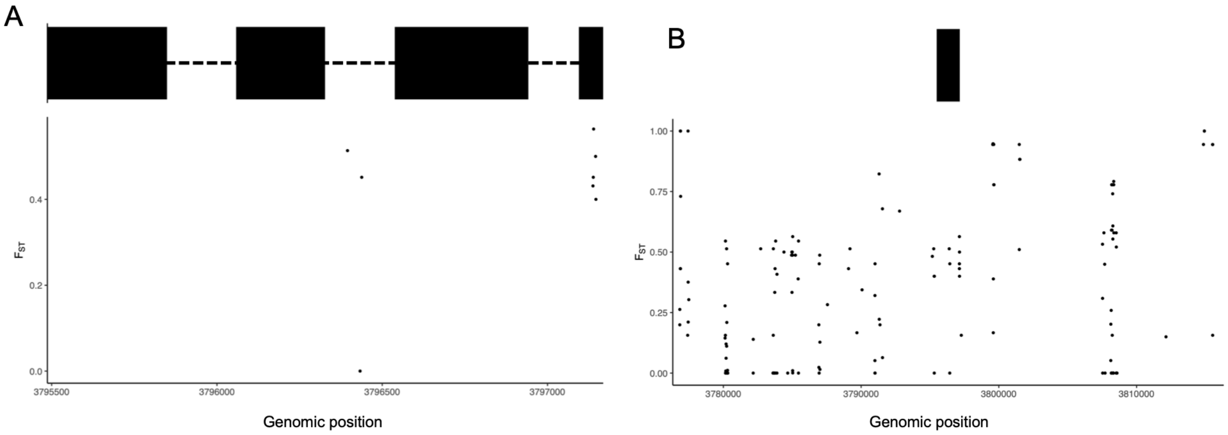


**Fig. S2.** SNP-level Fst A) within *GonOMT1*, where solid blocks represent exons and dotted lines representing introns, and B) 20 kb upstream and downstream of the gene (solid block).

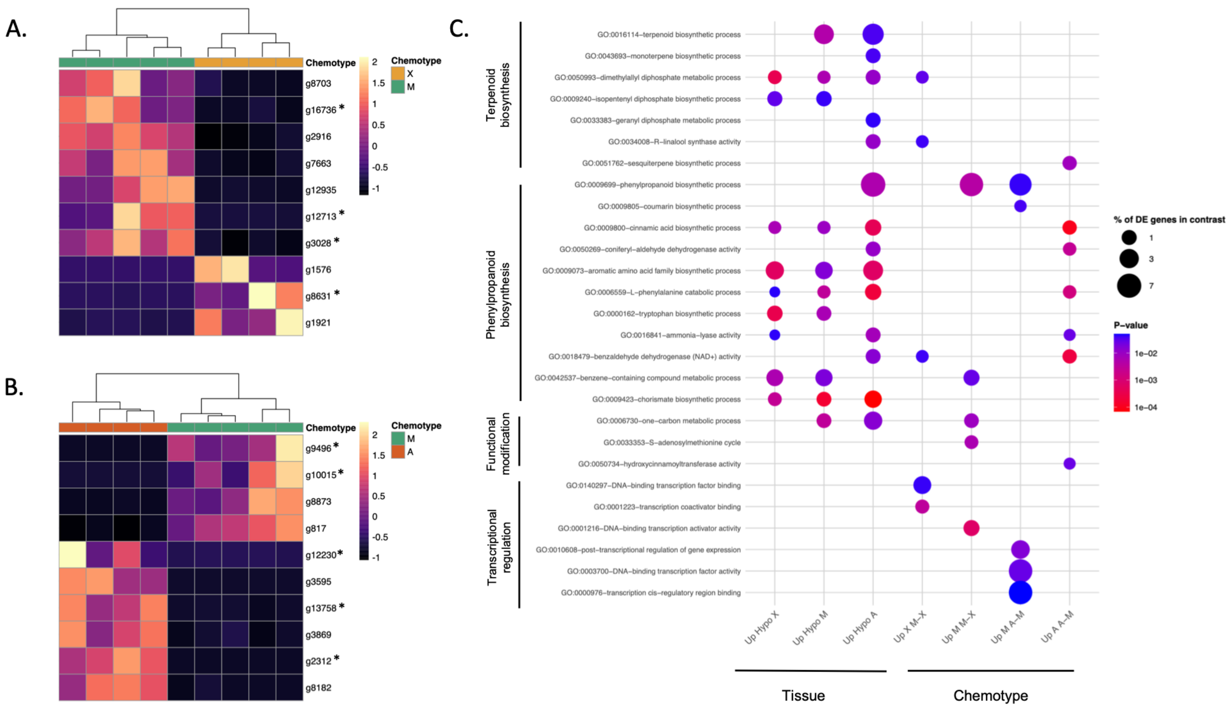


**Figure S3.** Major patterns of differential expression between contrasts and significantly enriched terms. A) and B) show heatmap of the top 10 most differentially expressed genes in the M-X and A-M contrasts, respectively. Genes marked with asterisks represent those that additionally exhibit significant upregulation in the hypochiles of the respective chemotypes. C) and D) show the most enriched terms associated with the DE genes of the two contrasts.

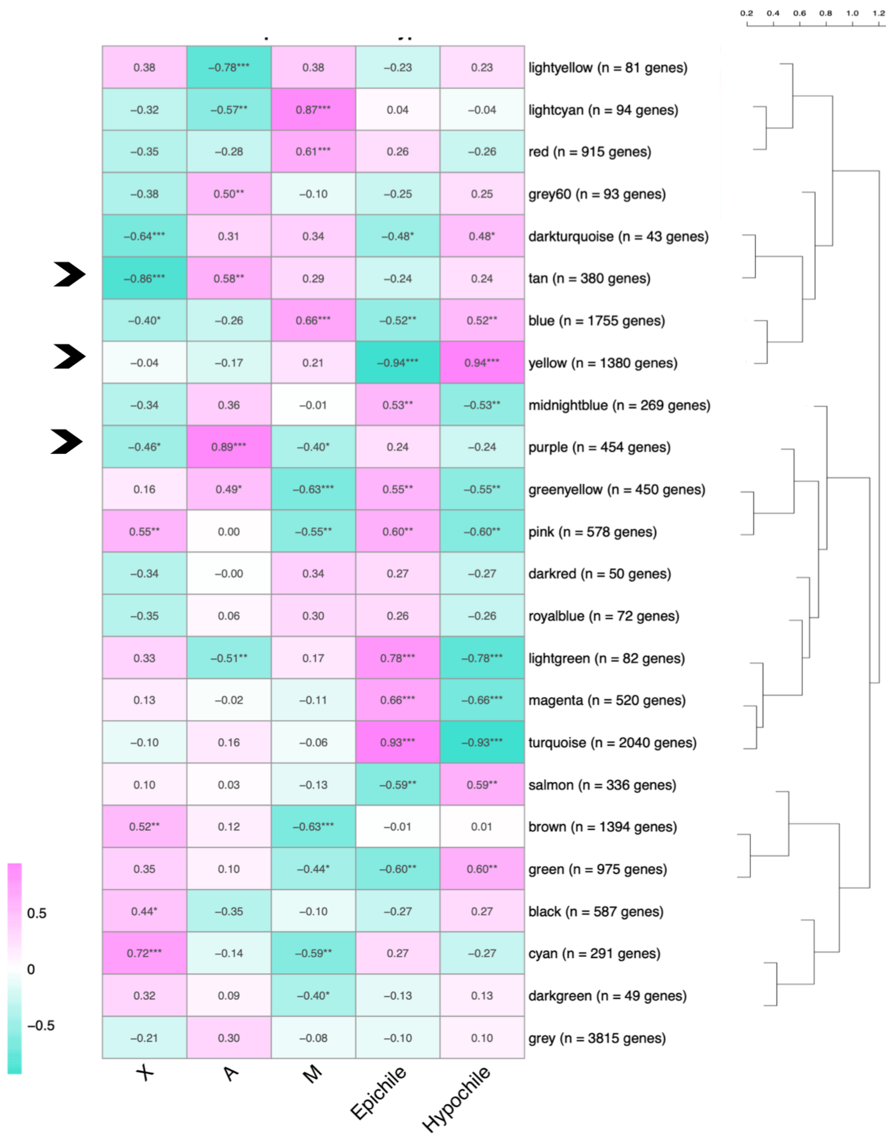


**Figure S4.** Results of WGCNA, with a heatmap of correlations between module eigengenes with chemotypes and with tissue. The dendrogram on the right shows the relationships between the different modules, with the axis above corresponding to dissimilarity in overall expression patterns. Values in the cells correspond to correlations between the category and module eigengene expression while * = significant at the 0.05 level, ** = significant at the 0.01 level, *** = significant at the 0.001 level. Arrows on the left side correspond to modules of interest mentioned in the main text, with “tan” corresponding to the “monolignol module”, “yellow” corresponding to the “hypochile-specific module”, and “purple” corresponding to the “aromatic modification module”.

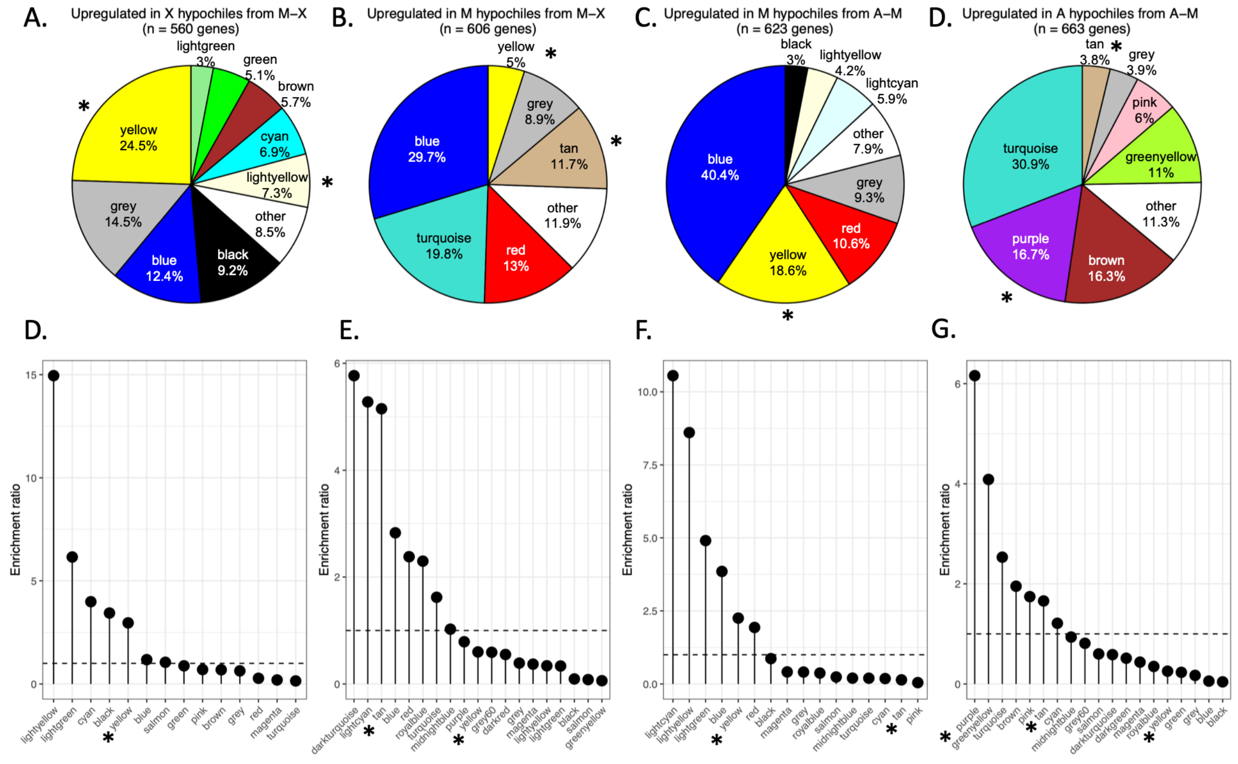


**Fig. S5.** Distributions of which modules differentially expressed genes from different contrasts were present in. Asterisks indicate modules mentioned in the text, with “purple” corresponding to the “aromatic-modification module”, “tan” corresponding to the “monolignol module”, and “lightyellow” corresponding to to the “hypochile-specific module”. A-D show pie charts of these distributions from upregulated genes from each chemotype in the two main contrasts (M-X and A-M) while E-H show ”enrichment” of modules for these respective categories, based on fold-increase from an expectation of equal distribution across modules (e.g. in D a value of 15 on the y-axis means that the lightyellow module had 15 times more genes than expected that were upregulated in X from the M-X contrast given its size). Dotted lines represent the expectation of 1-fold enrichment.

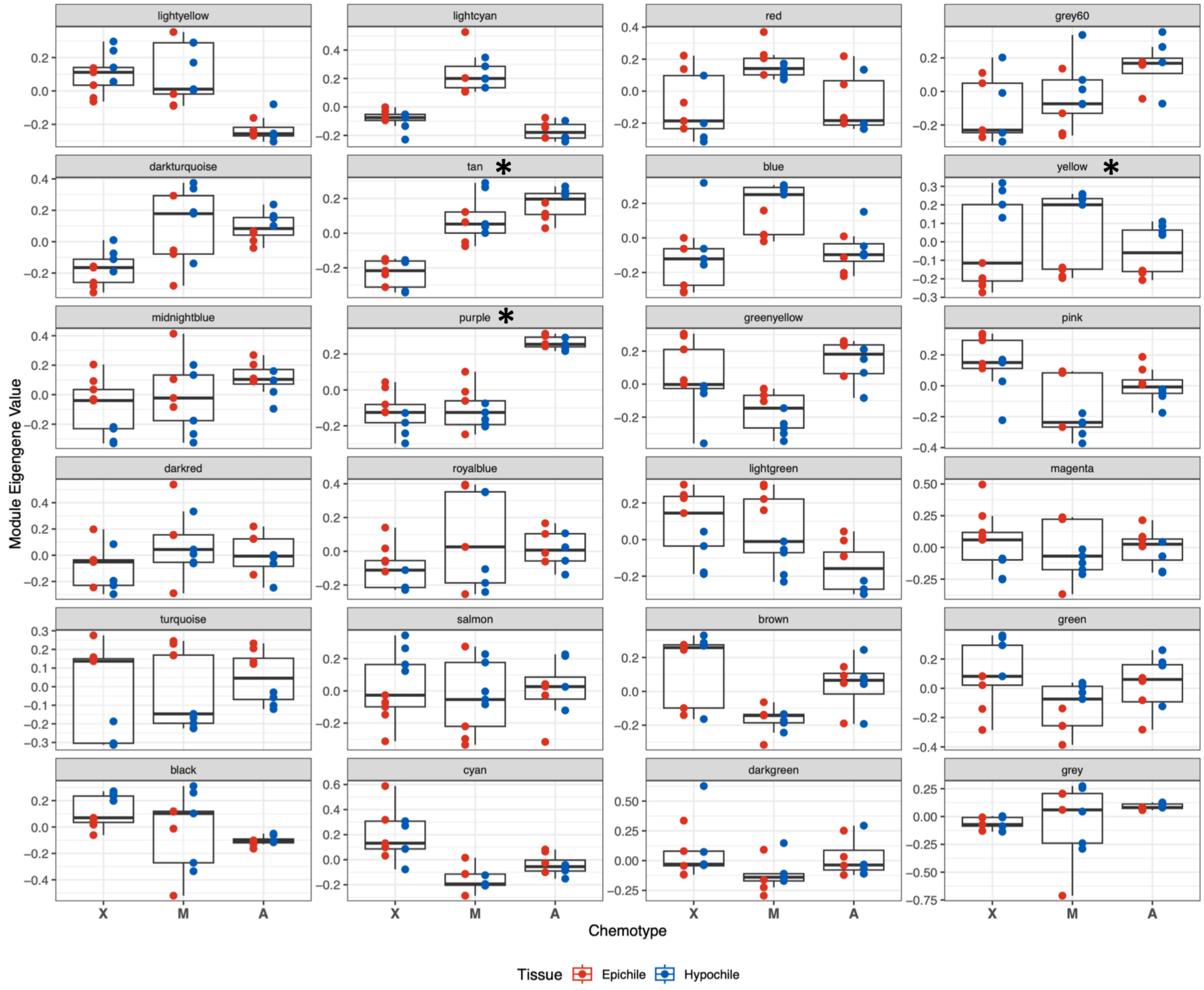


**Fig. S6.** Module eigengene values for all modules identified in WGCNA. Modules marked with asterisks correspond to the modules mentioned in the main text, with "tan" corresponding to the "monolignol module", "yellow" corresponding to the "hypochile-specific module", and "purple" corresponding to the "aromatic modification module".

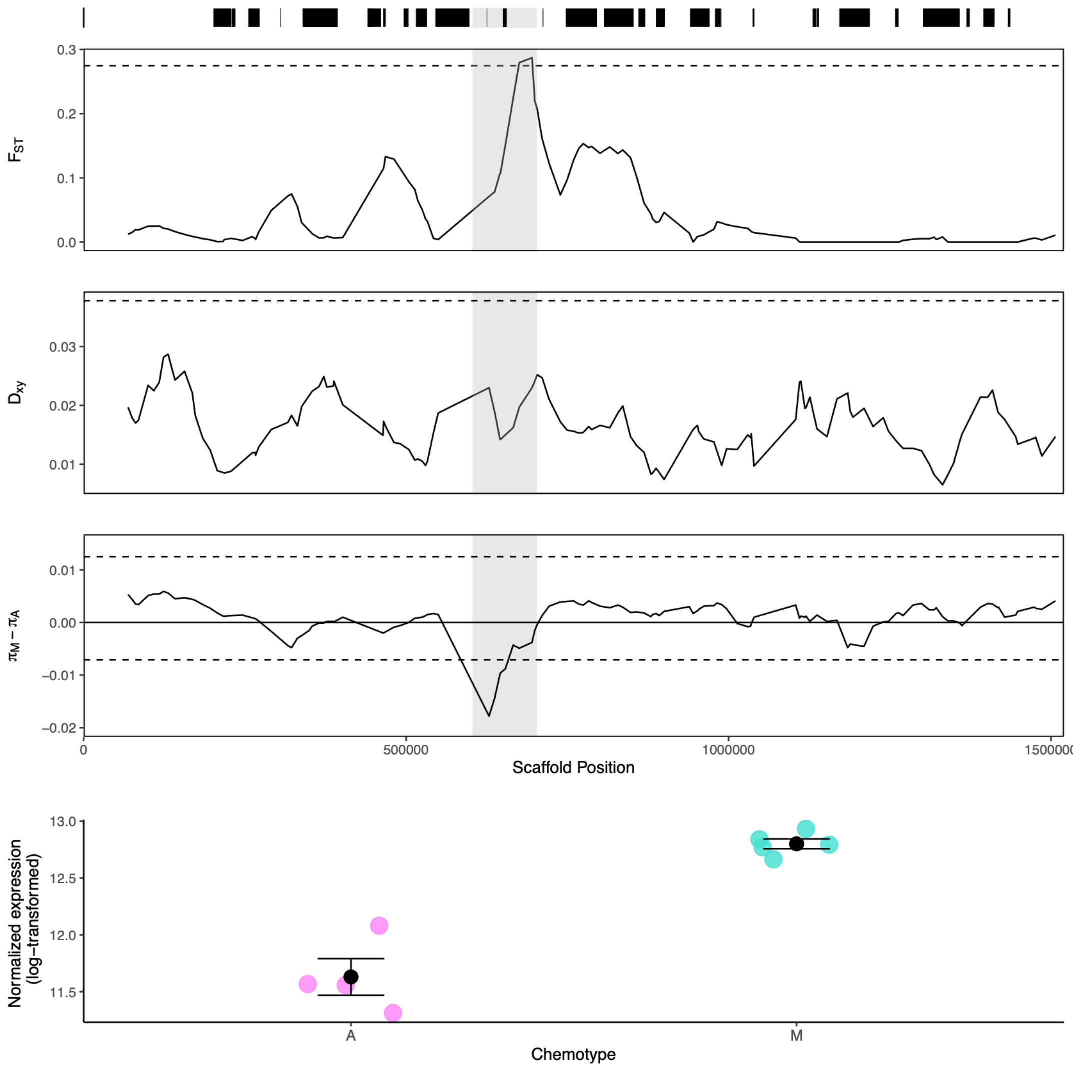


**Fig. S7.** Results of 50 kb sliding analysis for *GonDXPS1*. The black rectangles at the top represent genes, with the one surrounded by gray representing *GonDXPS1* and 50 kb around it. A. Sliding window estimates of Fst, with the dashed line representing the 99.9$^{th}$ percentile of 0.275. B. Sliding window estimates of Dxy, with the dashed line representing the 99$^{th}$ percentile of 0.0378. C. Sliding window estimates of heterozygosity of A subtracted from heterozygosity of M. The upper dotted line corresponds to the 99$^{th}$ percentile of this distribution (0.0125) while the lower dotted line corresponds to the 1$^{st}$ percentile of the distribution (-0.0078). D. Expression differences of *GonDXPS1* between the A and M chemotypes
